# Sensory stimulation triggers different spike responses in serotonin and dopamine neurons in the dorsal midbrain tegmentum

**DOI:** 10.64898/2026.06.11.731093

**Authors:** Péter Földi, Kinga Müller, Dániel Magyar, Gergő A. Nagy, Norbert Hájos

## Abstract

The dorsal midbrain tegmentum, including the dorsal raphe nucleus (DRN) and the ventrolateral periaqueductal gray (vlPAG), contains diverse neuronal populations. Within this region, serotonin (5-HT) and dopamine (DA) neurons are the principal monoaminergic cell types and exert widespread influence on brain circuits. While the role of 5-HT and DA neurons in sensory integration is well established, their stimulus-driven spiking activity remains incompletely characterized. Using silicon probe recordings in mice, we found that >57% of DRN/vlPAG neurons responded to foot shock and mechanical stimulation, whereas <15% showed changes in spiking activity following light or acoustic stimulation. At the population level, similar results were obtained using juxtacellular recordings, a method that allowed post hoc identification of 5-HT and DA neurons. Upon foot shock delivery, 5-HT neurons exhibited heterogeneous responses, including both excitation and inhibition, whereas DA neurons typically increased their firing rates. We found that DA neurons lacking vasoactive intestinal polypeptide (VIP) fired within the first second after foot shocks, while VIP-expressing DA neurons were most active later. Together, our results demonstrate that DRN/vlPAG neurons are most responsive to foot shock and mechanical stimuli. Moreover, 5-HT and DA neurons exhibit distinct patterns of activation following aversive inputs, suggesting that they play different roles in sensory information processing.

## Introduction

The ability to detect and respond to behaviorally relevant sensory stimuli, particularly those signaling potential threat or harm, is essential for survival. Aversive stimuli engage distributed brain networks that integrate sensory inputs with internal state to guide adaptive behavioral and physiological reactions ^1-4^. One of the key nodes in this survival network is the dorsal tegmentum of the midbrain, including the dorsal raphe nucleus (DRN) and the ventrolateral periaqueductal gray (vlPAG) ^3,5,6^. These neighboring structures have long been implicated in the modulation of pain, defensive behaviors, and arousal ^3,4,7-9^, yet the neuronal dynamics underlying their responses to sensory inputs remain incompletely understood.

The DRN and vlPAG contain heterogeneous populations of neurons, including glutamatergic, GABAergic neurons and monoaminergic cell types releasing serotonin (5-hydroxytryptamine, 5-HT) or dopamine (DA) ^10-13^. Among these, monoaminergic neurons give rise to widespread projections that innervate diverse cortical and subcortical regions, including the prefrontal cortex, basal ganglia, and amygdala ^10,11,14,15^. Through these monoaminergic projections, 5-HT and DA systems exert broad influence over cognitive, affective, and motivational processes ^1,16-18^. A large body of work has demonstrated that these neuromodulatory systems contribute to sensory processing, particularly in the context of aversive and rewarding stimuli ^1,16,17,19^. However, much of this knowledge derives from indirect measures or task-related activity monitoring Ca^2+^ transients ^1,16,19,20^, and relatively little is known about how these neurons respond at the level of spiking activity to well-defined sensory inputs. In particular, the extent to which monoaminergic neurons are differentially engaged by aversive versus non-aversive sensory inputs has not been systematically characterized.

Another unresolved question concerns the functional diversity within monoaminergic populations themselves. 5-HT neurons in the DRN have been shown to display heterogeneous responses to aversive stimuli ^15,16,20,21^, including both excitation and inhibition, suggesting that they may encode distinct aspects of sensory or emotional information. DA neurons, although less abundant in the dorsal midbrain tegmentum compared to other midbrain structures such as the ventral tegmental area ^13,18,19,22,23^, have also been implicated in processing salient and aversive signals ^1,18,19,22,24^. Recent work has further revealed molecular and functional heterogeneity among these DA neurons, including the presence of subpopulations defined by neuropeptide expression, such as vasoactive intestinal polypeptide (VIP) ^13,14,23,25^. How such diversity translates into differences in sensory stimulus-evoked spiking dynamics remains largely unknown.

Understanding how monoaminergic neurons in the dorsal midbrain tegmental neurons respond to different types of sensory stimuli is critical for elucidating their roles in integrating external inputs with internal states ^1,8,15,20^. In particular, comparing responses to aversive and neutral sensory modalities can reveal whether these circuits are specialized for processing behaviorally relevant stimuli or whether they broadly encode sensory information. Moreover, resolving the temporal dynamics of neuronal responses across distinct cell types may provide insight into how information is transformed and relayed to downstream target areas.

In the present study, we recorded sensory stimulus-triggered spiking activity of neurons in the DRN and vlPAG. Using silicon probe recordings in urethane-anesthetized mice, complemented by juxtacellular labeling to identify 5-HT and DA neurons, we systematically examined neuronal responses to foot shock, mechanical, visual, and auditory stimuli. Earlier studies showed that sensory stimulus-evoked responses are largely preserved under this type of anesthesia ^26-28^. Our results reveal that a substantial fraction of dorsal midbrain tegmental neurons is preferentially responsive to aversive foot shock and mechanical stimulation, while responses to visual and auditory inputs are comparatively rare. Furthermore, we find that 5-HT and DA neurons exhibit distinct response profiles to aversive stimuli, differing in both response type and temporal dynamics, indicating functional specialization within these neuromodulatory systems. Together, these findings provide new insight into how the dorsal midbrain tegmentum processes salient sensory information and contributes to adaptive behavioral responses.

## RESULTS

### Sensory inputs evoke diverse responses in DRN/vlPAG neurons

To assess neuronal activity in the DRN/vlPAG region in response to various sensory stimuli, we examined single-unit firing performing acute multi-channel silicon probe recordings in urethane-anesthetized male mice (N = 4; Fig. 1A). From 9 penetrations, we successfully identified 82 well-isolated single-units (for details, see Materials and Methods; Fig. 1B). Five types of sensory stimuli - foot shock, mechanical pressure, aversive noise, neutral tone, and bright light - were presented in a pseudorandom sequence with a 15-second inter-trial interval (Fig. 1A). Among these stimuli, foot shock and mechanical stimulation of the paw with a von Frey filament are considered aversive or strongly aversive ^29^, while the 18–20 kHz sweeping tone (aversive noise) evokes a mildly aversive response ^30^. In contrast, the pure 8 kHz tone and the 525 nm light are salient but neutral stimuli.

**Figure 1.**
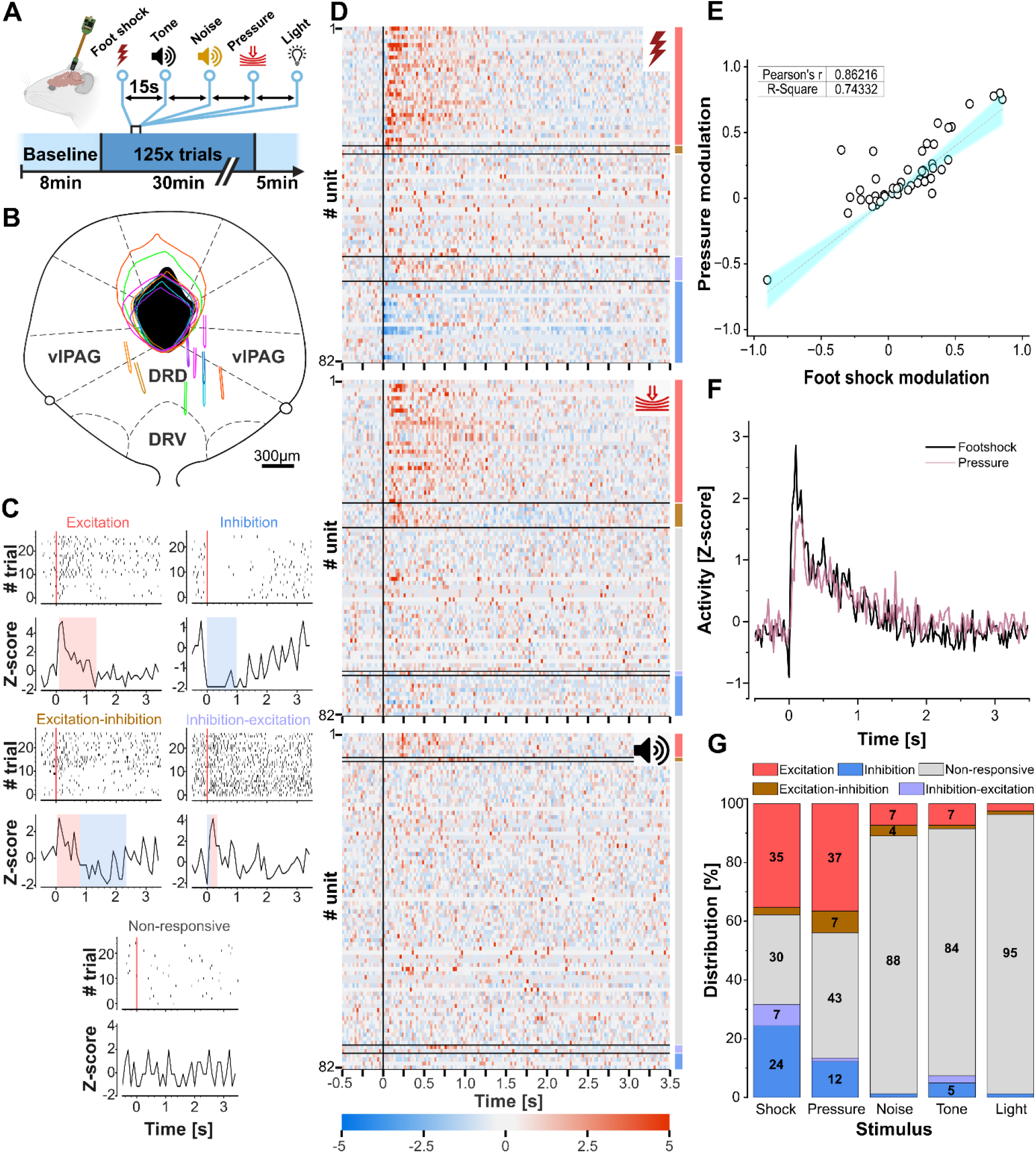
Characterization of single-unit responses to sensory stimuli in the DRN/vlPAG region. **A)** Schematic of experimental design in urethane-anesthetized mice. Stimuli were presented in a pseudorandom sequence (inter-trial interval: 15 s). **B)** Recovered position of silicon probe tracks relative to the cerebral aqueduct. The tracks and the corresponding aqueduct positions are in the same color (N = 9). Scale bar = 300 µm. **C)** Example peri-stimulus time histograms (PSTH) of single-units exhibiting different typical responses following foot shocks (Z-score bin size: 200 ms, dots represent single spikes). Blue and red shaded intervals show time windows where neurons were significantly excited and inhibited, respectively. **D)** Heatmaps showing the PSTHs of all single-units (N = 82) related to foot shocks (top panel), to mechanical (middle panel) and auditory stimulations (8 kHz, bottom panel). Units are sorted based on their response characteristics. Color coded bars adjacent to panels indicate the corresponding response type, (same color code as in G). The color scale of the heatmap represents Z-scores (bin size: 25 ms). Stimulation onset is marked at the zero-time point with a vertical black line. **E)** Correlation of modulation indexes for excitatory and inhibitory responses to foot shock or mechanical stimulation (N_units_= 47). Modulation indexes were calculated for 1 s post-, and 5 s pre-stimulus intervals. Solid line represents the linear regression fit; shaded area in blue indicates the 95% confidence band. **F)** Averaged population activity of excited neurons in response to aversive stimulation (N_units_= 29, bin size: 25 ms). **G)** Percentages of neurons with distinct response characteristics upon sensory stimuli.

To explore the temporal dynamics of spiking activity changes upon stimulation, we applied the Pruned Exact Linear Time (PELT) change point detection algorithm ^31^ to Z-score–normalized peri-stimulus firing rates of neurons. We identified three basic activity patterns: excitation, inhibition, and no change. Additionally, two biphasic patterns were observed: excitation followed by inhibition (excitation-inhibition) and inhibition followed by excitation (inhibition-excitation) (Fig. 1C). The majority of neurons significantly modulated their activity in response to foot shock (69.51%; Fig. 1D, top) and mechanical filament stimulation (57.32%; Fig. 1D, middle), with a strong correlation between the two response profiles (Pearson’s R = 0.862, p < 0.0001; Fig. 1E). Moreover, the spiking dynamics of neurons excited by foot shock and mechanical stimulation were indistinguishable (Figs. 1F, 2B). In contrast, only a small proportion of neurons responded to any of the other stimuli (<15% in all cases; Fig. 1G), suggesting that under anesthesia, predominantly aversive or strongly aversive stimulations drive significant activity changes in DRN/vlPAG neurons. The similarity in firing responses to foot shock and mechanical stimulation implies the involvement of shared afferent pathways conveying these signals to dorsal midbrain tegmental neurons.

### Aversive stimuli typically excite DA neurons, whereas spiking responses in 5-HT neurons are more diverse

Due to the cellular heterogeneity within the DRN/vlPAG region, our next goal was to identify the specific neuron types modulated by aversive stimuli. To achieve this, we performed single-cell recordings followed by Neurobiotin labeling using a juxtacellular recording configuration while delivering foot shocks or mechanical stimuli (Fig. 2A), both of which reliably evoked responses in most neurons within this dorsal tegmental region (Fig. 1G). In 80 urethane-anesthetized mice, we successfully visualized 140 neurons. Neurons that could not be unequivocally linked to their electrophysiological recordings or that were located outside the DRN/vlPAG were excluded from analysis (N = 13 mice, N = 20 neurons). Among the 120 juxtacellularly recorded neurons included in the dataset, we observed the same characteristic response types as those identified in silicon probe recordings (Fig. 2C, D). First, spiking responses to foot shocks strongly correlated with those elicited by mechanical stimulation (Pearson’s R= 0.719, p= 1.084 × 10^−4^, Fig. 2B). Second, all response types previously classified with silicon probe recordings were also identified in the juxtacellular dataset (Fig. 2C). Notably, the distribution of response types was comparable between the two methods (Figs. 1D, 2C). Specifically, the majority of neurons (39.17%) were excited by foot shock; 10.83% showed an excitation–inhibition pattern, 7.5% exhibited inhibition–excitation, and 13.33% displayed pure inhibition (Fig. 2D). A substantial fraction of neurons (29.17%) did not significantly alter their firing rate in response to foot shocks. Because the timing of foot shocks can be precisely controlled, we used this stimulation type in subsequent experiments.

**Figure 2.**
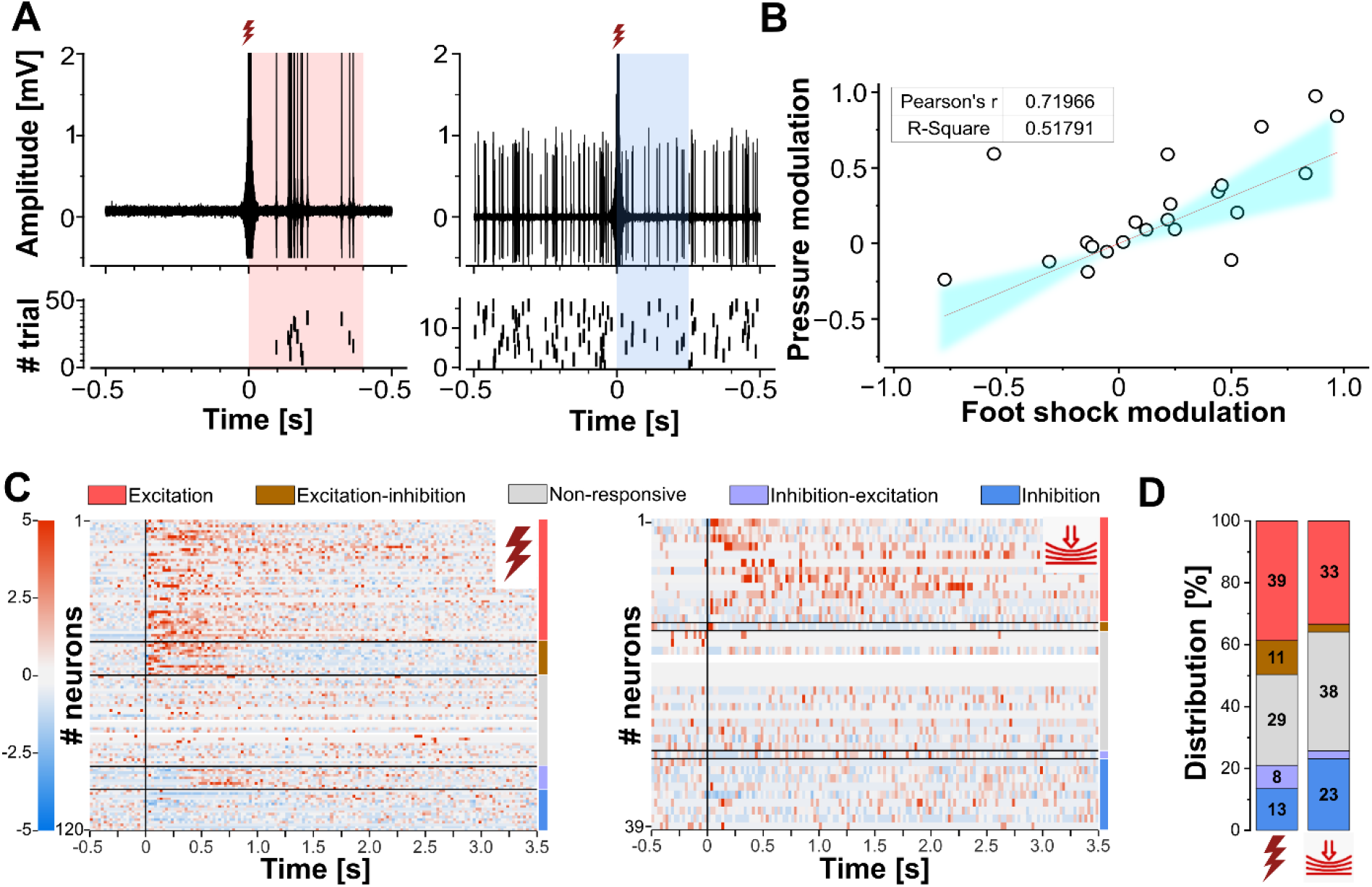
Population response of juxtacellularly recorded neurons to aversive stimuli in the DRN/vlPAG region. **A)** Example voltage traces from single neurons excited (left) or inhibited (right) by foot shock, recorded juxtacellularly. **B)** Correlation of modulation indexes for excitatory and inhibitory responses to foot shocks and mechanical stimulations (N_units_= 22). Modulation indexes were calculated for 1 s post-, and 5 s pre-stimulus intervals. Solid line represents the linear regression fit; shaded area in blue indicates the 95% confidence band. **C)** Heatmaps showing the PSTHs of juxtacellularly recorded neurons (N = 120) related to foot shocks (left panel) and to mechanical stimulation (N = 39 from 120 units, right panel). Spikes are sorted based on their response characteristics. Color coded bars on the right side of the panels indicate the corresponding response type. The color scale of the heatmap on the left represents Z-scores (bin size: 25 ms), (white color means no firing in the analyzed time period). Stimulation onset is marked at the zero time point with a vertical black line. **D)** Percentages of neurons with distinct response characteristics upon aversive stimulation. Color code is the same as in Fig. 1.

To confirm that juxtacellular and multi-channel recordings sampled comparable neuronal populations within the DRN/vlPAG, we related the stimulus-evoked response characteristics exclusively among neurons that exhibited a significant response to aversive stimulation. The distribution of peak Z-score timing was similar between the two recording methods (Mann–Whitney U test, *p* = 0.67; Fig. 3A), and there was no significant difference in response magnitude as measured by maximum Z-score change (Mann–Whitney U test, *p*_*exc*._ = 0.24 for excitatory, and *p*_*inh*._ = 0.35 for inhibitory response components; Fig. 3B). However, baseline firing rates prior to stimulation were significantly higher in neurons recorded with multi-channel silicon probes (Mann–Whitney U test, *p* = 1.11 × 10^−3^; Fig. 3F). Despite this difference, the response onset (*p*_*exc*._ = 0.26, *p*_*inh*._ = 0.44), offset (*p*_*exc*._ = 0.96, *p*_*inh*._ = 0.41), and duration (*p*_*exc*._ = 0.64, *p*_*inh*._ = 0.79) were statistically indistinguishable between recording modalities (Mann–Whitney U test; Fig. 3C-E). These findings suggest that both recording techniques sampled overlapping populations of functionally responsive neurons in the DRN/vlPAG.

**Figure 3.**
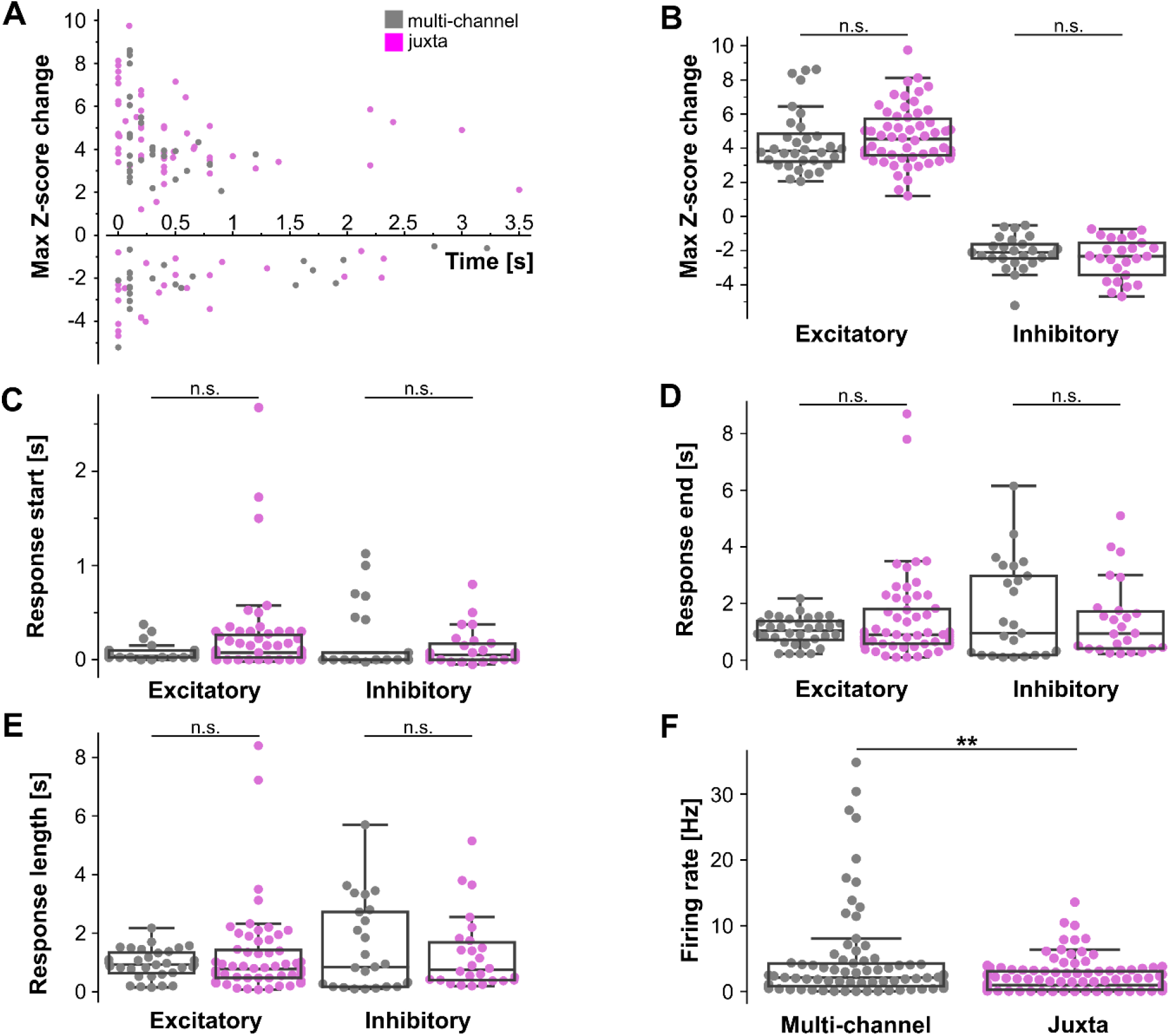
Comparison of population responses of neurons to foot shocks recorded with multi-channel silicon probes and juxtacellular recording technique. **A-B)** Maximal change in Z-scored firing rates after foot shock in all the analyzed neurons. **C-E)** Response onset, termination and total length of spiking in foot shock-responsive neurons (N_multi-channel_ = 57, N_juxta_ = 83). **F)** Basal firing rate of all recorded neurons (N_multi-channel_ = 82, N_juxta_ = 120). Color code: grey: multi-channel silicon probe recordings, magenta: juxtacellular recordings. Statistical differences were assessed using Mann-Whitney’s U-test (see text for details).

To identify the neurochemical content of recorded neurons, we performed immunocytochemical staining on Neurobiotin-filled juxtacellularly recorded neurons. Specifically, we investigated their immunoreactivity against 5-HT, tyrosine hydroxylase (TH), and VIP, which are abundantly expressed in the DRN/vlPAG ^10^. TH is the rate-limiting enzyme in DA synthesis ^32^ and serves as a marker for DA neurons in this region, while VIP is expressed in a subset of TH-expressing cells ^13,33^. Using multicolor confocal microscopy, we found that 38 neurons (31.67%) were immunopositive for 5-HT, 7 neurons (5.83%) were exclusively immunoreactive for TH (TH+VIP-), and 12 neurons (10.00%) co-expressed TH and VIP (TH+VIP+, Fig. 4B-D). The remaining 63 neurons (52.50%) did not show detectable expression of any of the three markers and were classified as neurochemically unidentified. Due to the possibility that this group may include multiple cell types - some potentially expressing low levels of the tested markers - they were excluded from subsequent statistical analyses. Consistent with prior reports ^11,34^, 5-HT neurons were located predominantly along the midline and lateral wings of the DRN. DA neurons identified by TH immunostaining were more broadly distributed throughout the DRN/vlPAG region ^25,33^, while TH+/VIP+ neurons were primarily located near the ventral part of the cerebral aqueduct, consistent with earlier observations ^13,35^ (Fig. 4A).

**Figure 4.**
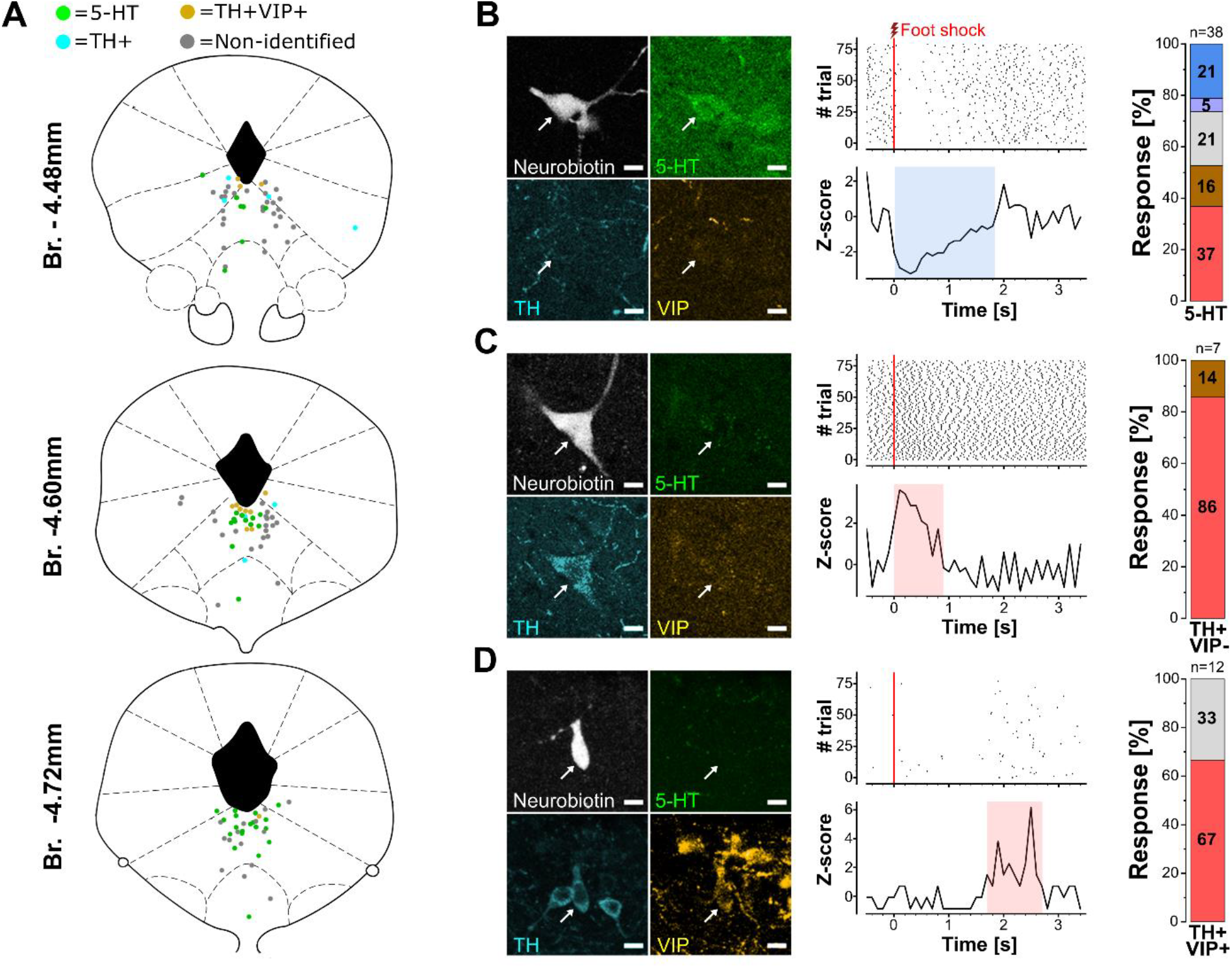
Foot shock-evoked responses in juxtacellularly recorded and neurochemically identified neurons in the DRN/vlPAG. **A)** Localization of Neurobiotin-filled neurons in the DRN/vlPAG region. Colors indicate neurochemical marker positivity (green: 5-HT, turquoise: TH+VIP-, yellow: TH+VIP+, grey: immunonegative for 5-HT, TH and VIP). Br. = Bregma levels. **B-D)** Fluorescence visualization of Neurobiotin-filled example neurons and the results of immunostaining for 5-HT, TH and VIP (left panels, arrows show the labelled neurons, scale bar: 10 µm) alongside their corresponding PSTHs upon foot shocks (middle panel) and distributions of the characteristic responses of neurons with the same neurochemical markers (right panel, color code is the same as in Fig. 1G). Red vertical lines in PSTHs indicated stimulus onset, whereas the highlighted periods where neuronal activity increased (red) or decreased (blue).

We assessed foot shock-evoked responses in neurons with distinct neurochemical content and found that 5-HT cells (N=38) exhibited a diverse range of characteristic responses (excitation: 36.84%, excitation-inhibition: 15.79%, inhibition-excitation: 5.26%, inhibition: 21.05%, non-responsive: 21.05%, Fig. 4B). In contrast, DA neurons predominantly displayed excitatory responses to foot shocks (in TH+VIP-neurons (N=7), excitation: 85.71%, excitation-inhibition: 14.29%, Fig. 4C; in TH+VIP+ neurons (N=12), excitation: 66.67%, non-responsive: 33.33%, Fig. 4D).

To examine the temporal dynamics of neuronal responses across neurochemically identified cell types, we focused on neurons that exhibited excitation or excitation–inhibition activity patterns following foot shocks (Fig. 5A-B). Analysis of excitatory responses only revealed that TH+VIP+ neurons displayed significantly delayed response onsets (0.944 ± 0.913 s, data are presented as mean ± standard deviation) compared to TH+VIP− (0.046 ± 0.057 s) and 5-HT neurons (0.101 ± 0.152 s; Kruskal–Wallis test, H = 14.63, *p* = 6.664 × 10^−4^; Fig. 5C). Similarly, response offsets were significantly prolonged in TH+VIP+ neurons (2.363 ± 0.848 s) relative to TH+VIP− (1.025 ± 1.139 s) and 5-HT cells (0.907 ± 0.681 s; Kruskal–Wallis test, H = 11.29, *p* = 3.533 × 10^−3^; Fig. 5D). In contrast, there were no significant differences in the duration of excitatory responses (Kruskal–Wallis test, H = 5.19, *p* = 0.074; Fig. 5E) or in their peak normalized amplitude (Kruskal–Wallis test, H = 2.24, *p* = 0.327; Fig. 5F) across cell types. Neurons exhibiting inhibitory or inhibition–excitation responses were excluded from this analysis, as such patterns were observed exclusively in 5-HT neurons following foot shocks.

**Figure 5.**
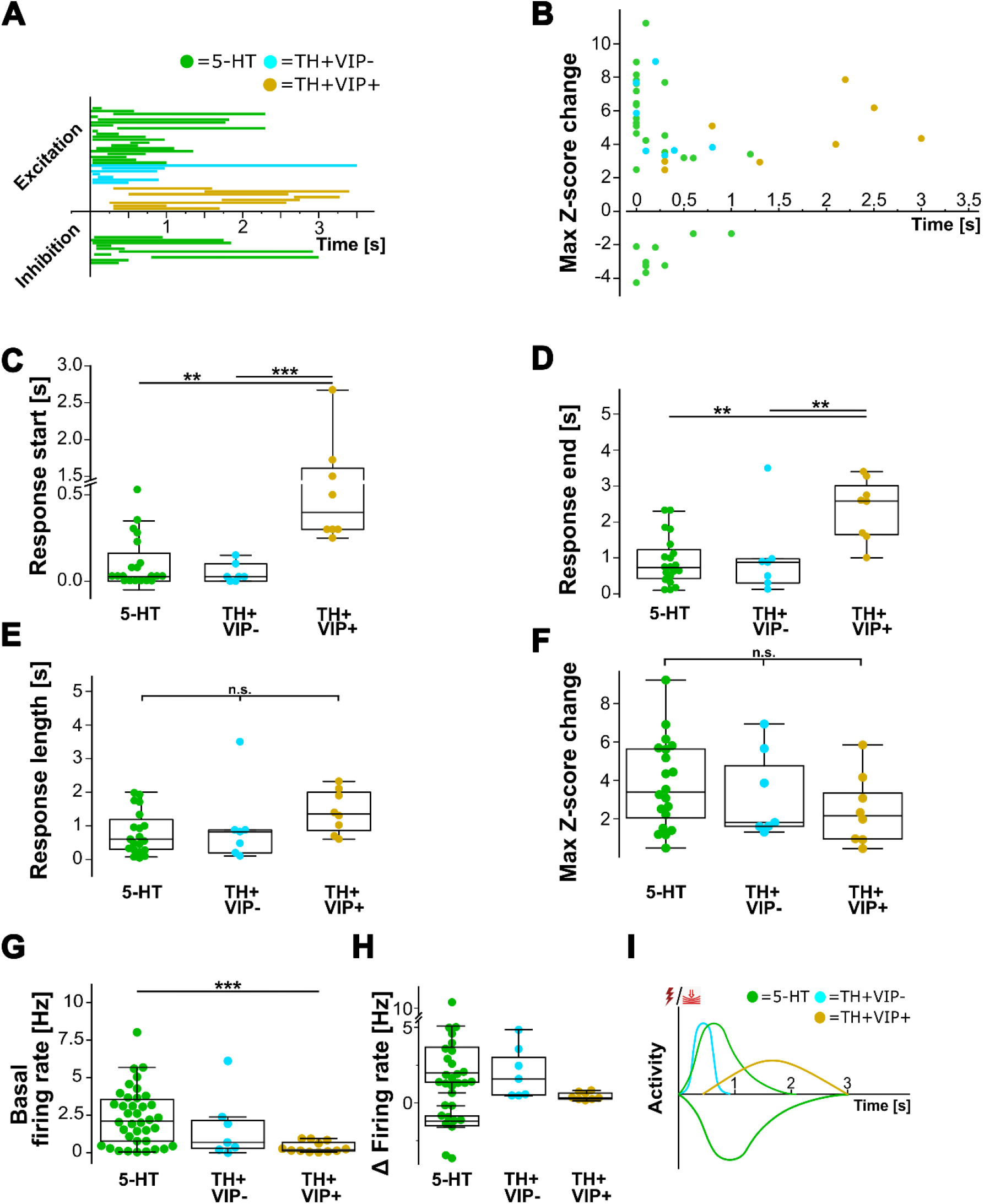
Aversive stimulation-triggered changes in firing in neurochemically identified neurons in the DRN/vlPAG. **A)** Response time windows for individual neurons upon foot shock stimulation. Time windows were determined based on change point detection of Z-scored firing rates (PELT searching method). **B)** Maximal change in Z-scored firing rates after foot shocks in identified neurons (N_5-HT_ = 20, N_TH+VIP-_ = 7, N_TH+VIP+_ = 8). **C-F)** Onset, termination, total length and maximal magnitude of excitatory responses only (N_5-HT_ = 20, N_TH+VIP-_ = 7, N_TH+VIP+_ = 8). **G)** Basal firing rates of juxtacellularly recorded neurons. **H)** Firing rate change upon foot shock stimulation of responsive neurons. **I)** Schematic summary of activity change in neurochemically identified neurons upon aversive stimulation. Color code: green: 5-HT, turquoise: TH+VIP-, yellow: TH+VIP+. Z-score bin size for A, C-E panels: 25 ms; B, F panels: 100 ms. Statistical differences were assessed using Kruskal-Wallis ANOVA, followed by post hoc Dunn’s test. Significant pairwise differences are indicated (* p < 0.05; ** p < 0.01; *** p < 0.001).

### Intrinsic firing properties of neurochemically identified neurons

Finally, we examined the baseline firing characteristics of neurochemically identified neurons prior to stimulus delivery. A comparison of basal firing rates (BFRs) across cell types revealed a significant difference (Kruskal–Wallis test, H = 13.45, *p* = 1.2 × 10^−3^; Fig. 5G). TH+VIP+ neurons exhibited a significantly lower mean BFR (0.38 ± 0.34 Hz) compared to 5-HT neurons (2.38 ± 1.86 Hz; Dunn’s post hoc test, *p* = 2.82 × 10^−4^). While TH+VIP+ neurons also showed a trend toward lower BFRs relative to TH+VIP-neurons (1.67 ± 1.99 Hz), this difference was not statistically significant (Dunn’s test, *p* = 0.1366). Given the large variability in 5-HT neuron BFRs, we further assessed whether firing rates differed among 5-HT neurons based on their response patterns. However, no significant differences were observed across response subtypes (Kruskal–Wallis test, H = 7.16, *p* = 0.13; Table 1).

**Table 1.**
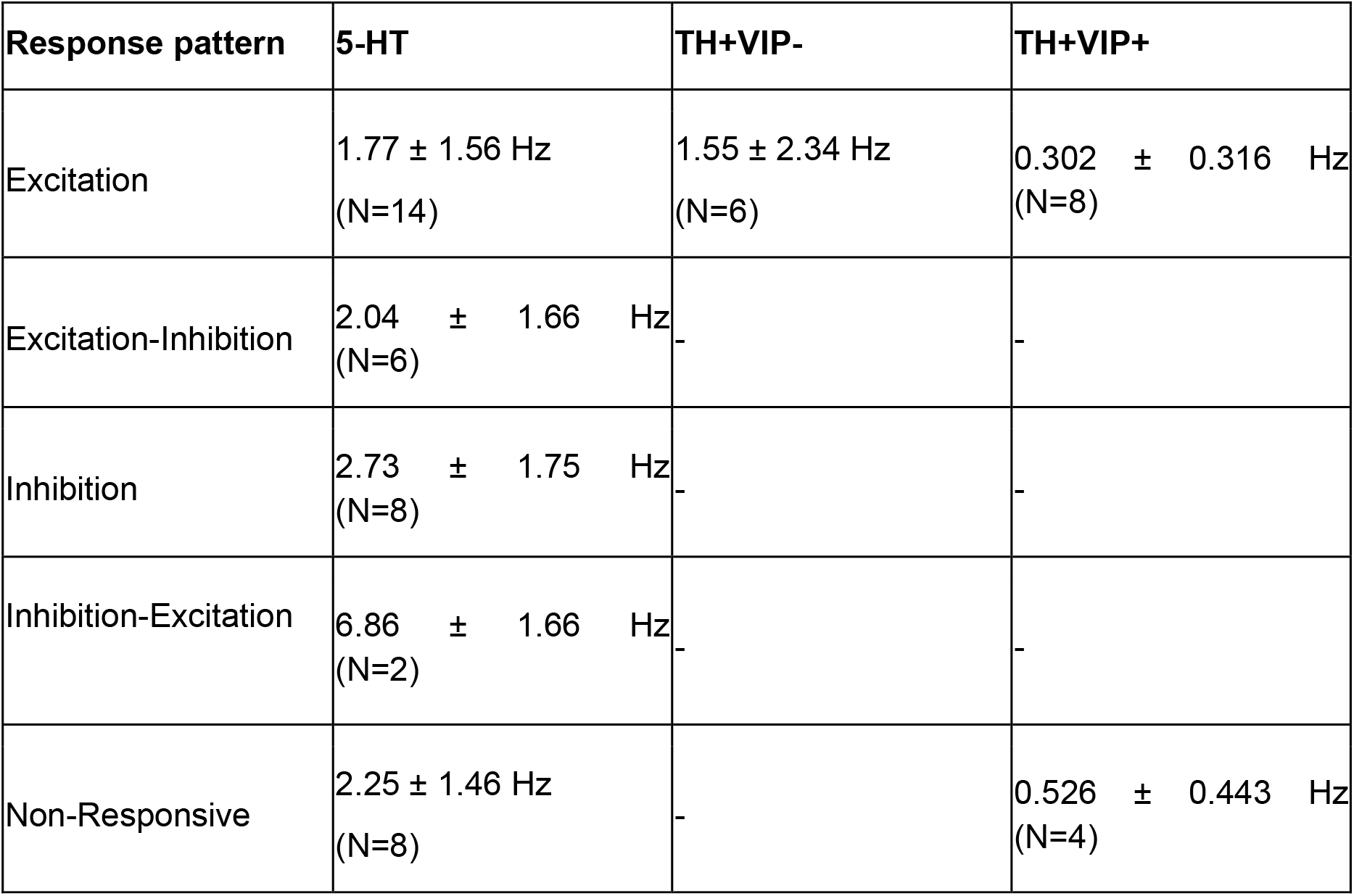
Mean basal firing rates for neurochemically identified DRN/vlPAG neurons categorized by response pattern. Values are reported as mean ± SEM only when N>1.

Previous studies have revealed that in this region, neurons may display two characteristic firing patterns: regular spiking and burst firing ^36-39^. In our recordings, we identified a subset of neurons (N = 6) with a low coefficient of variation (CV) of inter-spike intervals (ISIs; CV < 0.33), indicative of regular, pacemaker-like firing. Consistent with previous reports ^36,40^, the majority of these neurons (N = 4) expressed 5-HT. Additionally, approximately 10% of neurons (N = 12) exhibited burst-like firing patterns, defined by the presence of doublets or triplets and at least 10 ISIs shorter than 20 ms ^37,38^. This burst firing spiking phenotype was observed in all monoaminergic neurons studied, albeit rarely. These observations indicate that regular and burst firing characteristics do not separate monoaminergic neuron types in the dorsal midbrain tegmentum.

## Discussion

The main findings of our study are as follows. 1) Foot shock and mechanical stimulation altered spiking activity in most neurons (70% and 57%, respectively) in the DRN and vlPAG. Spike responses to foot shock and pressure delivery showed a linear relationship. Only a small fraction of DRN/vlPAG neurons (<15%) responded to acoustic and light stimulation. 2) Spiking activity in the majority of 5-HT neurons (79%) changed in response to aversive stimulation; a larger fraction showed increased firing (53%), whereas a smaller fraction exhibited inhibition (26%). 3) DA neurons were typically excited by shocks (>67%). Foot shock-evoked spike dynamics in DA neurons correlated with their VIP content. These results indicate that sensory stimuli mostly excite a population of neurons in the DRN and vlPAG, with similar spiking responses to both aversive foot shock and mechanical stimulation. In addition, monoaminergic neurons in this region exhibit distinct patterns of excitation following aversive inputs, suggesting that they play different roles in sensory information processing.

Our findings provide a systematic characterization of how diverse sensory stimuli affect neuronal activity in the dorsal midbrain tegmentum, specifically in the DRN and vlPAG, and extend previous work by directly comparing responses across multiple modalities within the same experimental conditions. Earlier studies have demonstrated that neurons in the DRN and vlPAG respond robustly to aversive stimulation, including those inducing feelings of pain, defensive behavior, and arousal ^9,22,24,41^. In particular, electrophysiological recordings in the PAG have shown strong activation by noxious stimuli such as foot shock, pinch, or thermal inputs ^9,41,42^, whereas responses to non-aversive modalities are less consistently reported ^1,16,20,22^. Our observations are in agreement with these findings, showing that a large fraction of neurons in the DRN and vlPAG respond to both foot shock and mechanical stimulation, while only a small proportion are modulated by visual or auditory inputs. Importantly, our data further demonstrate that responses to foot shock and mechanical pressure are highly correlated at the single neuron level, suggesting that these dorsal tegmental circuits may encode a common dimension of somatosensory salience rather than modality-specific features. These observations extend previous work by indicating that dorsal midbrain tegmental neurons are broadly tuned to behaviorally relevant somatosensory signals ^9,41-43^, with limited engagement by distal sensory modalities such as visual and auditory stimuli under the conditions tested.

Our results also refine the current understanding of how 5-HT neurons process sensory information. Previous studies using electrophysiology, Ca^2+^ imaging, and optogenetic approaches have reported heterogeneous responses in DRN 5-HT neurons to aversive stimuli, including both excitation and inhibition ^15,16,20,21^. These mixed responses have been interpreted as reflecting functional diversity within the 5-HT system, with different subpopulations potentially encoding distinct aspects of sensory, emotional, or motivational states. Our findings are consistent with this view, as we observed both increases and decreases in spiking among 5-HT neurons following aversive stimulation. However, our data further indicate that excitatory responses predominate, suggesting that the net effect of aversive input on 5-HT output may be biased toward increased activity. This aligns with studies showing that subsets of 5-HT neurons are activated by stressors and noxious stimuli and contribute to behavioral responses such as anxiety-like behavior and behavioral inhibition ^15,21,44,45^. At the same time, the presence of inhibitory responses in a subset of neurons supports the idea that 5-HT neurons are not homogeneous and may contain functionally distinct neuron types that are differentially engaged by sensory inputs ^15,16,20,21^. By directly linking spiking responses to defined sensory modalities, our results provide additional evidence that 5-HT neurons participate in encoding salient somatosensory events while maintaining internal heterogeneity that could support diverse downstream functions.

In contrast to 5-HT neurons, DA neurons in the dorsal midbrain tegmentum have been less extensively characterized, particularly in relation to sensory-evoked spiking ^1,22,24,43^. Studies examining DA neurons within or near the DRN/vlPAG region have suggested that these cells can be activated by stress and aversive events ^1,22,24,43^, but detailed analyses of their firing dynamics have not been conducted. Our findings provide clear evidence that DA neurons in the dorsal midbrain tegmentum are robustly excited by aversive stimulation. Moreover, we show that the temporal dynamics of these responses are not uniform but instead depend on the presence of VIP. DA neurons lacking VIP respond rapidly within the first second after stimulation, whereas VIP-expressing DA neurons exhibit delayed activation. This temporal segregation suggests that distinct subpopulations of DA neurons may encode different phases of information processing, such as immediate detection versus sustained evaluation of salient stimuli. Alternatively, they may provide a prolonged effect in their downstream regions, the bed nucleus stria terminalis and central nucleus of the amygdala ^13,14^. Our observations add to the growing body of literature emphasizing heterogeneity within DA systems ^13,14,25,46^ and suggests that dorsal tegmental DA neurons contribute to sensory processing in a manner that is both functionally and temporally specialized.

Taken together, our findings show that the DRN and vlPAG can integrate behaviorally relevant sensory information, particularly signals related to potential threat or bodily perturbation. Rather than serving as general-purpose sensory relays, neurons in these regions appear to preferentially encode somatosensory stimuli that are likely to require immediate behavioral or physiological responses ^9,22,24,41,42^. The convergence of responses to foot shock and mechanical stimulation, combined with the relative insensitivity to visual and auditory inputs, supports the idea that the dorsal midbrain tegmentum is tuned to proximal, high-salience signals ^1,8,22,41,42^. Importantly, 5-HT neurons exhibit heterogeneous response patterns that may support flexible modulation of downstream circuits ^15,16,20,21^. In contrast, DA neurons in the dorsal midbrain tegmentum display more uniform excitation with subtype-specific temporal dynamics, potentially endorsing differentially salient signal processing ^1,18,22,24,43^. These differences suggest that 5-HT and DA systems are not redundant but instead provide parallel pathways through which sensory information can influence brain-wide activity. By revealing how distinct neuronal populations in the dorsal midbrain tegmentum respond to different classes of sensory stimuli, our study helps understanding how these circuits contribute to adaptive behavior ^2,8,18,43^ and how their dysfunction may underlie disorders associated with altered processing of behaviorally relevant stimuli, such as anxiety and stress-related conditions ^9,24,43,47,48^.

## Materials and Methods

### Animals

In all experiments, adult C57Bl/6 mice were used (mean age: 128±48.7 days for juxtacellular and 183+24.7 days for silicon probe recordings; (N = 38 females and N = 42 males for juxtacellular and N = 4 males for silicon probe recordings). Animals were group-housed under a 12-h light/dark cycle with food and water available *ad libitum*. All experimental procedures were approved by the Committee of the Scientific Ethics of Animal Research operated under the National Food Chain Safety Office (NÉBIH) (22.1/360/3/2011) and by the Animal Welfare Body (MÁB) of HUN-REN Institute of Experimental Medicine. All procedures were conducted in accordance with the relevant national legislation (1998 XXVIII. section 243/1998, renewed in 40/2013.) and the institutional ethical guidelines. Experiments complied with the European Convention for the Protection of Vertebrate Animals Used for Experimental and Other Scientific Purposes (Directive 86/609/EEC, as revised by Directive 2010/63/EU). All efforts were made to minimize animal suffering and to reduce the number of animals used. The study also adhered to the ARRIVE guidelines.

### Surgical procedure

Mice were anaesthetized with isoflurane (2% v/v, CP-Pharma GmbH) carried by O_2_ (at 0.4 L/min) and then mounted in a standard stereotaxic instrument (Kopf Instruments®). Before any incision lidocaine (10%) was applied locally and corneal gel (Vidisic, Bausch & Lomb) was used to protect the eyes. The animal’s body temperature was continuously maintained at approximately 37°C.

A small craniotomy was conducted (anteroposterior, AP -5.3 mm; mediolateral, ML 0.0 mm for silicon probe recordings and AP -4.7 mm, ML 0.0 mm for juxtacellular recordings) with great caution due to exposure to the sinus. To avoid compromising audition, a lightweight head plate (Neurotar) was affixed to the skull horizontally using denture acrylic (Paladur). A pool was constructed around the craniotomy from denture acrylic to provide additional protection and prevent brain tissue dehydration. Following the surgical procedure, urethane (∼0.8 g/kg i.p., in 10% solution, Sigma-Aldrich, Cat# U2500) was administered to the mice and the recording session commenced a few minutes later.

### Stimulations and electrophysiological recordings

#### Multi-channel silicon probe recordings

Mice were positioned into the setup (HeadFix, Neurotar) by firmly securing the head plate into the holder, ensuring stable head fixation. For single-unit recordings a 2 × 32 channel silicon probe with pre-installed optic fiber (H10 ASSY-77, Cambridge NeuroTech) was acutely inserted into the brain with 20° tilt along the axial axis. The shanks of the silicon probe were immersed in an ethanol-dissolved lipophilic dye, DiI (Sigma-Aldrich) for histological reconstruction of electrode tracks after the recordings. As a ground electrode, an Ag-AgCl electrode was placed under the animal’s scalp. The silicon probe was positioned and gradually advanced at a rate of 1-2 µm/second using a micromanipulator (SM7, Luigs & Neumann GmbH) until the target depth was reached (2800-3400 µm from dura mater) ^49^. To prevent dehydration of the brain surface, saline solution was applied every 10-15 min. Electrical signals were recorded using an Intan RHD-2000 system (Intan Technologies), connected via Neuronexus AO32 × 2 adaptor to two 32-channel amplifier boards (RHD2132, Intan Technologies). Data were digitalized and recorded with a sampling rate of 30 kHz and the resolution of 16 bits (OpenEphys, v0.6.6.) ^50^.

During a recording session 5 different types of external stimuli were introduced: foot shock (3 ms, 1 mA), mechanical stimulus (30-35 gram-force, for 500 ms), neutral tone (8 kHz, ∼90 dB, for 500 ms), mildly-aversive noise (100 ms long tone sweeps from 18 to 20 kHz, 90 dB, for 500 ms ^30^, and bright light (525 nm, 20 µW, for 500 ms). For foot shock stimulation, two metal conductors were attached to both hind paws of the animals, that transmitted a weak current emitted by the stimulator (BioStim, SuperTech Ltd), causing a mild electric shock. Before every set of stimulation, a highly conductive gel (ECG gel, Rextra) was applied between the metal conductors and the paws. Mechanical stimulation was provided by a von Frey-like monofilament which was fabricated based on ^29^ operated by a small servo motor hitting the paw of the animal. For auditory stimulations two loudspeakers (5 W, FRS-5x, Visaton) were connected to a stereo audio-amplifier (TDA7297 chip-based). Visual stimulation was delivered using two LEDs positioned 10 cm from the animal’s eye. All stimulatory devices were connected and controlled via an Arduino Uno R3, which was synchronized by TTL signals with the CED 1401 A/D converter (delay: <60 ns, Cambridge Electronic Design). Stimuli were introduced in a pseudo-random order with the inter-stimulation time of 15 s. A complete protocol contained 125 trials (25 of each stimulus). After an approximately 8-minute-long recording of the baseline firing activity, the stimulation protocol was executed, while continuous recordings of the electrical signals were maintained, followed by slow retraction of the silicon probe. In some cases, another recording was initiated with an alteration of 500 µm on the ML or 250 µm along the AP axis.

#### Juxtacellular recordings

For juxtacellular recordings, a glass micropipette was fabricated from borosilicate capillaries (Sutter Instruments, BF150-86-10, impedance measured by bridge-balance: 18-30 MΩ) and filled with 3% Neurobiotin solution (Vector Labs, dissolved in 0.5 M NaCl). Signals were transmitted from a headstage (CV-7B, Axon Instruments), amplified (MultiClamp 700B, Molecular Devices), filtered (50-60 Hz elimination, Hum Bug Noise Eliminators), digitalized at 20 kHz (CED 1401, Cambridge Electronic Design) and recorded with Spike2 (Cambridge Electronic Design). An Ag/AgCl electrode was used as the reference. Since approaching the target area (2900-3400 μm from the dura) requires vertically penetrating through the sinus, slow advancement (1-2 μm/s) of micropipette was required, which prevented bleeding and caused the brain tissue to retain its original shape, providing more stable recordings.

For a subset of juxtacellular recordings (N = 39) the stimulation protocol included the same 5 stimuli described above presented in the same pseudo-random order. For the remaining recordings (N = 81), the stimuli protocol consisted of 20-80 mild electric foot shocks (3 ms, ∼1 mA), delivered alternately between the paws, with a consistent inter-stimulation interval of 15 s.

For juxtacellular labelling ^51^ with Neurobiotin, positive, rectangular nano currents (200 ms with 400 ms intervals) were delivered, gradually increasing the current from 100 pA until the cell showed current-modulated firing. This state was maintained for as long as possible, up to a maximum of 20 min. Depending on the anesthesia and the quality of the recording, the entire recording session was repeated with a minimum of 150 μm change in the AP or DV axis to be able to distinguish the labelled neurons.

### Histology

After recordings, animals – under the long-lasting anesthetic effect of urethane administered before the recordings – were injected intraperitoneally with another dose of urethane (∼1.6 g/kg, in 20% solution, Sigma-Aldrich, Cat# U2500) to establish deep anesthesia. After confirming the complete loss of nociceptive reflexes mice were perfused directly through the circulatory system with 0.9 % saline for 2 min followed by 120-150 ml, 4 % paraformaldehyde (Sigma-Aldrich, Cat# 158127) fixative in phosphate buffer [0.1 M phosphate buffer (PB), pH=7.4]. The brain was immersed in 0.1M PB or fixative for a minimum of 12 h, depending on the quality of fixation.

To confirm the location of silicon probes, coronal sections (150 µm) were cut using a vibratome (VT1000S, Leica Biosystems) and washed for 3 × 15 min in 0.1 M PB. After mounting the sections, the DiI labelled silicon probe tracks were analysed by epifluorescent microscopy. Recordings outside the DRN/vlPAG region were discarded from the dataset.

Brains containing juxtacellularly labelled neurons were cut to 40 µm-thick coronal sections using a vibratome (VT1000 S, Leica Biosystems) and washed (3 × 15 min in 0.1 M PB and in tris-based buffered saline (TBS) for another 3 × 15 min). For immunostaining against Neurobiotin, sections were incubated overnight in a solution containing Cy3-conjugated Streptavidin (SA, 1:20,000 in TBS, Sigma-Aldrich, Cat# S6402). Sections were incubated in a mixture of primary antibodies developed against 5-HT (rat anti-5-HT, 1:500, in 1:2 glycerol, Millipore, Cat# MAB352, RRID:AB_11213564), TH (chicken anti-TH, 1:2000, in 1:2 glycerol, ImmunoStar, Cat# 213 106, RRID:AB_2782977), and VIP (rabbit anti-VIP, 1:2000, in 1:10 0.1M PB, ImmunoStar, Cat# 20077, RRID:AB_572270) for 24 h at room temperature and for 4 days at 4°C. Then, sections were washed (3 × 20 min in 0.1 M PB) and stained with Alexa488-conjugated donkey anti-rat (1:500, Molecular Probes, Cat# A-21208, RRID:AB_2535794), Alexa647conjugated donkey anti-rabbit (1:500, in 1:2 glycerol, Jackson, Cat# 711-605-152, RRID:AB_2492288), DyL405-conjugated donkey anti-chicken (1:500, in 1:2 glycerol, Jackson, Cat# 703-475-155, RRID:AB_2340373), and 1% NDS containing solution for 3 h. After washing (3 × 15 min in 0.1 M PB), sections were mounted onto slides (ProLong− Diamond Antifade Mountant) and analysed using a confocal microscope (Nikon Ni C2 Confocal Microscope, Z-stacks with 1 μm step size).

### Data analysis

#### Analysis of multi-channel silicon probe recordings

Raw electrophysiological signals were introduced to Kilosort 2.5 ^52^ algorithm for spike detection and sorting with default parameters (without drift correction). Sorting was followed by manual curation using Phy2 (https://github.com/cortex-lab/phy; https://zenodo.org/records/3367782). The following parameters were considered upon quality check of unit isolation: auto correlogram for clean refractory period, cluster shape given by the 1^st^ and 2^nd^ principal component, waveform consistency and amplitude stability, cross-correlogram between similar units and waveform dissimilarity between units. Clusters with non-sufficient quality were excluded from further analysis. Timestamps of stimulations and unit activity were manually verified by reloading the raw data with labels into NeuroScope2 and identifying stimulation-driven noise detected as spikes. For visual inspection of unit responses and general features the data were loaded into CellExplorer ^53^.

Timestamps of spikes and stimuli were imported into a custom-written Python analysis pipeline, where the data was pre-processed and analyzed. To detect the onset and offset of neuronal activity alteration upon stimulation consequently, we applied a mean-drift-based multiple change point detection method called Pruned Exact Linear Time (PELT) algorithm ^31^ with fine-tuned parameters on Z-scored neuronal firing data (bin size: 25 ms). This method segments the peri-stimulus neuronal activity as a time series by minimizing a cost function with an added penalty term to control the number of detected change points, thus reducing overfitting. Due to the huge variability in firing rate – and thus the Z-scores – between neurons, manual curation of change points was indispensable. Within the framework of manual curation, we removed any change points outside the 5 s post-stimulation period; any segments with < 200 ms length outside 1^st^ second of post-stimulation period. Additionally, we adjusted the penalty parameter of the PELT algorithm, for cases with especially low firing rate, to identify potential changes in the activity. The spikes present between the change points (i.e. putative response time windows) were counted and were compared with the pre-stimulus spike count in multiple corresponding time windows. Statistically significant changes (Wilcoxon signed-ranked test with discarding all zero-differences, p < 0.05) were accepted as stimulus-induced changes, which served as a basis of clustering neurons into response classes. The BFR was calculated for the whole pre-stimulation time window.

For comparing the effects of different stimuli, we calculated modulation index and ratio using the same formula as implemented in CellExplorer ^53^ but restricted to time windows with significant change in the activity upon stimulation. For correlating these variables between stimuli, the data was grouped from all multi-channel recording sessions.

#### Analysis of labelled neurons and juxtacellular recordings

Cell-type classifications of juxtacellularly labelled neurons were based on the visual inspection of confocal images. In cases when the labelled cell could not correspond confidently with an electrophysiological recording (e.g., labelled cells were close to each other) or if cell-type identity was questionable, the data were excluded from further analysis. Labelled cells outside the DRN/vlPAG region were also excluded from the dataset. Spikes were identified in a threshold-based manner (Spike2) with additional manual curation. Timestamps of spikes and stimuli were imported into the same custom-written Python pipeline as introduced above. In short, Z-scored data (bin size: 25 ms) were introduced to change point detection with the PELT searching method. Change points were manually curated as described above. The BFR was calculated as the average spike count over the 100-s period preceding the first stimulus. Neurons that could not be unequivocally identified neurochemically were excluded from statistical comparisons, since this group may involve multiple types of neurons. To identify burst-firing neurons, we applied the criterion that at least 10 of all inter-spike intervals (ISIs) must be less than 20 ms. For comparisons of response latencies and intensities, non-parametric tests such as Kruskal-Wallis *H*-test with Dunn’s comparison (for N_groups_ > 2) or two-sided Mann-Whitney *U*-test (for N_groups_ = 2) were applied.

## DATA AVAILABILITY

The datasets generated and analyzed during the current study are available from the corresponding author on request.

## ACKNOWLEDGEMENTS

We acknowledge financial support from the HUN-REN Hungarian Research Network, Hungarian Brain Research Program (2017-1.2.1-NKP-2017-00002) and National Research, Development and Innovation Office (K131893) to N.H.. The authors are grateful to László Barna and Pál Vági, the Nikon Microscopy Center at the Institute of Experimental Medicine, Nikon Austria GmbH, and Auro-Science Consulting, Ltd., for kindly providing microscopy support.

## FUNDINGS

HUN-REN Hungarian Research Network

National Research, Development and Innovation Office (K131893)

Hungarian Brain Research Program (2017-1.2.1-NKP-2017-00002)

## AUTHOR CONTRIBUTIONS

P.F. carried out the *in vivo* electrophysiological recordings and their analysis, contributed substantially to immunocytochemical staining and analysis, and wrote the first draft of the manuscript. K.M. carried out the immunocytochemical investigations, contributed to their analysis, and reviewed the first draft. D.M. supervised the *in vivo* electrophysiological recordings and provided feedback on the initial draft. G.A.N. supervised the *in vivo* electrophysiological recordings and provided feedback on the initial draft. N.H. initiated and supervised the project, reviewed, and revised the initial draft, and approved the final version of the manuscript.

